# Augmenting Speech Comprehension using Rapid Frequency Tagging

**DOI:** 10.64898/2026.07.08.737094

**Authors:** Charlie Reynolds, Yali Pan, Ana Pesquita, Ole Jensen, Katrien Segaert, Hyojin Park

## Abstract

Speech comprehension under adverse listening conditions benefits from visual speech, but whether visual input can be modulated to improve listening remains unclear. Here we used rapid frequency tagging, a non-invasive sensory stimulation technique, in which the visual tag at 55 Hz over the mouth region of a talking face was amplitude-modulated by the envelope of either the task-relevant or task-irrelevant speech stream. When the visual modulation followed the task-relevant speech envelope, comprehension improved relative to task-irrelevant modulation and to an unmodulated tagging control. MEG responses showed enhanced 55 Hz visual tagging, altered 40 Hz auditory tagging and a non-linear 15 Hz intermodulation. Individual comprehension gains were associated with intermodulation responses in left inferior frontal cortex, linking behavioural benefit to audiovisual interaction. These findings show that task-relevant visual stimulation can improve comprehension by strengthening audiovisual interaction, establishing amplitude-modulated frequency tagging as a tool for probing and supporting comprehension in challenging environments.

## Introduction

Listeners must navigate speech-in-noise across everyday contexts, such as crowded restaurants, busy offices and social gatherings^1^. Speech comprehension in these environments depends not only on analysing the acoustic signal, but also on integrating visual information from the speaker’s face and body. Visual cues, such as lip movements and co-speech gestures, can guide attention^2^ and support comprehension^3,4^ by providing temporal and articulatory cues to the acoustic properties of speech, thereby aiding phoneme discrimination^5^. Furthermore, listeners can use visual information during naturalistic communication to generate predictions about upcoming speech, facilitating efficient language processing^6^. However, the ability to select and understand speech in noisy environments varies substantially across individuals. Listeners with neurodevelopmental conditions^7^, age-related hearing loss^8^ or cochlear implants^9^ often face particular challenges in understanding speech-in-noise. Identifying how speech cues can be manipulated to support comprehension in challenging listening conditions is therefore important both theoretically and practically. Here, we investigate whether speech comprehension in difficult listening contexts can be enhanced by manipulating speech-relevant visual cues.

Most approaches to improving speech-in-noise comprehension have focused primarily on enhancing auditory processing. However, growing evidence suggests that enhancing visual speech information may offer a promising an alternative means of supporting speech perception. Increasing the visual salience of the speaker’s mouth has been shown to improve speech intelligibility under degraded listening conditions^10^, suggesting that targeted enhancement of visual speech cues can support audiovisual perception. Here we asked whether speech comprehension can be improved by using visual input to deliver behaviourally relevant temporal information from the auditory speech signal, rather than simply by making visual speech cues more salient.

The speech envelope provides a natural candidate for such manipulation. It reflects slow amplitude fluctuations over time that convey rhythmic and prosodic information in speech^11,12^ and plays a crucial role in speech understanding and cortical speech tracking^13–15^, particularly in noisy environments^16,17^. Previous research has shown that speech comprehension was improved by expanding or enhancing the acoustic speech envelope cues^18,19^. The speech envelope has also been used as a form of visual rhythmic stimulation. For example, comprehension improved when participants viewed a visual stimulus whose size fluctuated congruently, rather than incongruently, with the auditory speech envelope^20^. Such findings provide evidence that visual presentation reflecting speech envelope dynamics can support speech perception in noise. However, such approaches have typically relied on simplified visual displays. We here ask whether auditory envelope information can be embedded in naturalistic visual input, such as the speaker’s mouth region, to enhance comprehension during continuous, naturalistic audiovisual speech.

Frequency tagging offers a powerful approach to address this question while also tracking neural responses to sensory information. In this approach, sensory stimuli are presented at specific temporal rates, eliciting steady-state neural responses at the corresponding frequencies^21–23^. This allows concurrently presented signals to be distinguished in the neural responses, as demonstrated for overlapping visual features tagged at different frequencies^24^. Conventional visual frequency tagging, however, has often used lower stimulation frequencies, making the flicker perceptible and placing the evoked response within frequency ranges that also show endogenous oscillatory and resonance activity^25^. The visual cortex can nevertheless follow periodic stimulation across a broad frequency range, with steady-state responses observed up to at least 90 Hz and frequency-dependent resonance effects reported around 10, 20, 40 and 80 Hz^26^.

Rapid Invisible Frequency Tagging (RIFT) builds on this capacity by applying visual stimulation above the perceptual flicker-fusion threshold^27^. The high-frequency visual tag can therefore be rendered imperceptible while leaving lower-frequency neural activity available for analysis, making RIFT particularly suitable for naturalistic paradigms^28^. RIFT has been used to investigate attention^25^, statistical learning^29^, speech planning^30^, natural reading^31^ and audiovisual speech processing^32^. The frequency tagging framework can also be used to measure interactions between auditory and visual signals. When the two modalities are tagged at different frequencies, modality-specific responses can be measured separately at their respective tagging frequencies. If the signals interact nonlinearly, additional responses can emerge at intermodulation frequencies corresponding to sums and differences of the original tags^33,34^. Because these frequencies are not present in either input alone, intermodulation responses provide a frequency-specific index of nonlinear audiovisual interaction. Drijvers et al.^32^ combined auditory tagging at 61 Hz with visual RIFT at 68 Hz and identified a 7 Hz difference-frequency response that was strongest when clear speech was accompanied by a congruent gesture. This response was localised to left inferior frontal and temporal regions, which have independently been implicated in integrating semantic information conveyed by speech and gesture^35^. Thus, modality-specific tagging and intermodulation responses provide complementary measures of auditory processing, visual processing and their nonlinear interaction.

In the present study, we move beyond this approach by using RIFT not only to measure audiovisual processing, but also to manipulate naturalistic visual speech. Specifically, we applied a 55 Hz visual tag to the speaker’s lip region and modulated its amplitude with the auditory speech envelope. This allowed the temporal structure of speech to be embedded directly into the visual speech signal. Crucially, the modulation was derived either from the speech stream that participants were instructed to attend or from the competing speech stream they were instructed to ignore. This design allowed us to test whether visual envelope modulation improves comprehension because it carries behaviourally relevant speech information, rather than because rhythmic visual stimulation is present per se. Specifically, we tagged auditory speech at 40 Hz and applied a 55 Hz visual tag to the speaker’s lip region and amplitude-modulated this signal with the auditory speech envelope, allowing behaviourally relevant speech to be embedded directly into visual input. This design tests whether speech comprehension can be improved when visual speech carries the temporal structure of the attended auditory stream, rather than merely providing a salient visual cue.

We hypothesised that embedding the attended speech envelope into visual speech would improve speech comprehension. At the neural level, we expected task-relevant modulation to strengthen visual tagging responses and alter auditory tagging responses, reflecting a shift in how auditory and visual speech signals are weighted during comprehension. Finally, we predicted that task-relevant modulation would enhance intermodulation responses at the 15 Hz difference frequency, providing a frequency-specific index of nonlinear interaction between auditory and visual speech signals.

## Results

To investigate whether visual rapid invisible frequency tagging (RIFT) modulated by the task-relevant speech envelope influences speech comprehension, participants completed a speech comprehension task under different tagging conditions. Figure 1 illustrates the experimental paradigm, including the dichotic speech task, the visual and auditory tagging frequencies, and the amplitude modulation conditions (Figure 1a,b). Representative coherence spectra showed clear peaks at the visual and auditory tagging frequencies, confirming robust steady-state responses at 55 Hz and 40 Hz, respectively (Figure 1c,d). The corresponding sensor topographies showed that visual tagging coherence was strongest over occipital sensors, whereas auditory tagging coherence was strongest over temporal sensors, consistent with the expected sensory origins of the two responses.

**Fig. 1.**
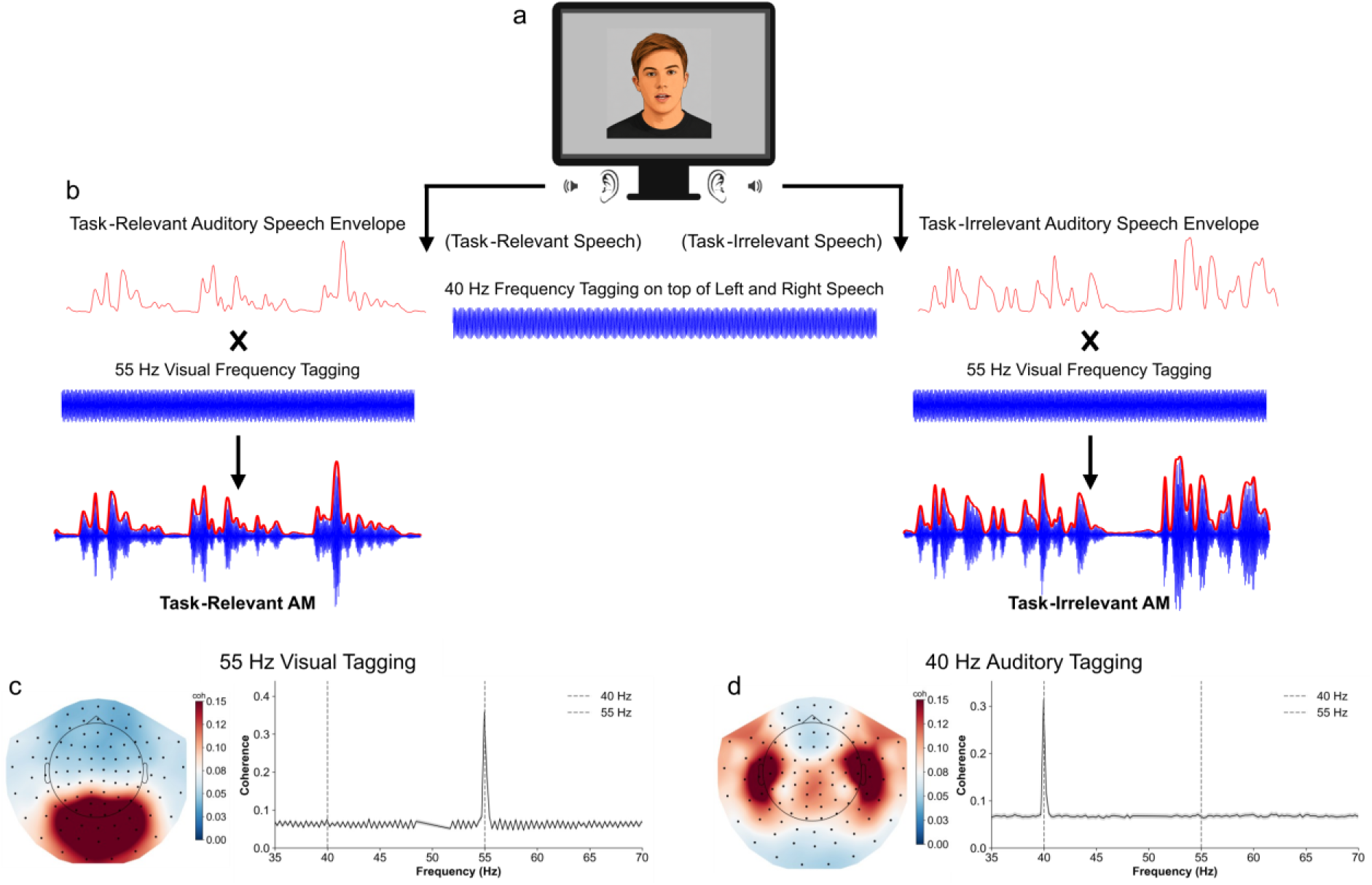
Experimental paradigm, stimulus construction, and representative frequency-tagging responses. **(a)** Participants viewed a centrally presented talking face while listening to two simultaneously presented speech streams delivered dichotically. For illustration purposes, the speaker is shown schematically. Auditory speech was frequency-tagged at 40 Hz, and the visual speech signal (lip region luminance) was frequency-tagged at 55 Hz. Participants were instructed to attend to one speech stream while ignoring the other. **(b)** Example construction of the frequency-tagged auditory and visual stimuli. Red traces show speech amplitude envelopes, and blue traces show sinusoidal tagging signals. The envelopes were multiplied by the carrier signals (visual frequency tagging at 55 Hz) to generate amplitude-modulated (AM) tagged stimuli. Depending on the experimental condition, modulation could be derived from the task-relevant (attended) or task-irrelevant (unattended) speech stream. **(c-d)** Representative coherence spectra and sensor topographies illustrating the steady-state evoked responses to the visual (55 Hz) and auditory (40 Hz) tagging signals. Coherence spectra show pronounced peaks at the respective tagging frequencies (dashed lines), while topographical maps reveal the spatial distribution of coherence across MEG sensors. The 55 Hz visual tag produced the strongest coherence over occipital sensors, whereas the 40 Hz auditory tag produced the strongest coherence over temporal sensors.

Specifically, we compared conditions in which amplitude modulation (AM) was either behaviourally relevant to the task (Task-Relevant AM), present but unrelated to the task (Task-Irrelevant AM), or absent while visual frequency tagging was still applied (Pure Tagging) (Figure 1). This design allowed us to dissociate the effect of behaviourally relevant envelope-based visual modulation from the effect of visual frequency tagging alone. Behavioural performance across these conditions is shown in Figure 2. The main text focuses on the three conditions central to the study question; results from the remaining experimental conditions and the full details of statistical comparisons are reported in the Supplementary Information.

**Fig. 2.**
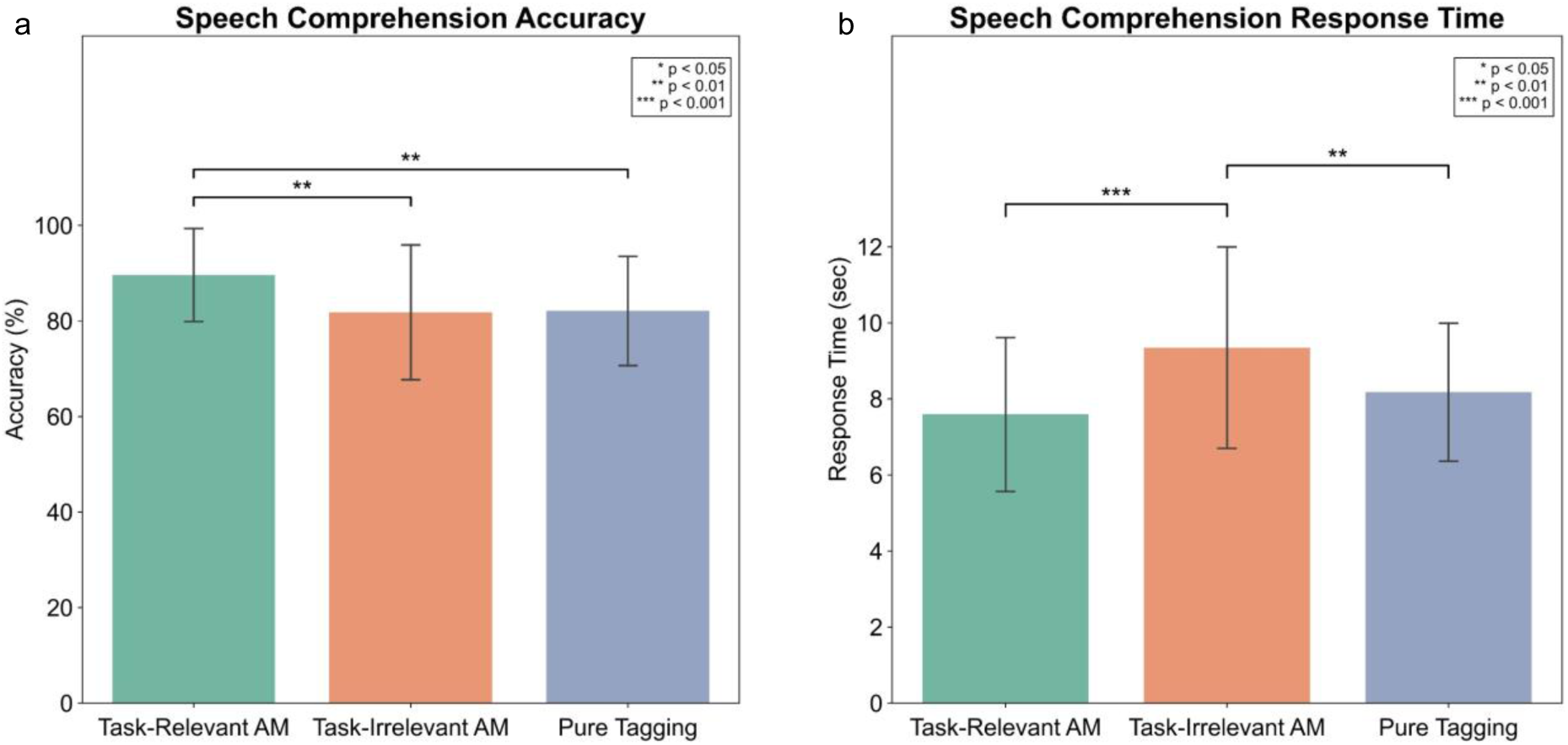
Speech comprehension results. **(a)** Mean speech comprehension accuracy across conditions. Accuracy was significantly higher in the Task-Relevant AM condition compared to both the Task-Irrelevant AM and Pure Tagging conditions. **(b)** Mean response times across conditions. Response times were significantly slower in the Task-Irrelevant AM condition compared to both Task-Relevant AM and Pure Tagging conditions. Error bars represent ±1 standard error of the mean. Statistical significance was assessed using one-way repeated-measures ANOVAs followed by Bonferroni-corrected paired-samples t-tests.

### Task-Relevant Amplitude Modulation Improves Speech Comprehension

We first asked whether behaviourally relevant amplitude modulation (AM) of the 55 Hz flicker of the visual input, guided by the auditory speech envelope, improves speech comprehension performance. A one-way repeated-measures ANOVA conducted across all five experimental conditions revealed significant differences in comprehension accuracy (F(4,156) = 4.82, p < 0.001; Supplementary Figure S1a). Given our a priori focus on the role of behaviourally relevant AM, subsequent Bonferroni-corrected paired-samples t-tests focused on the three focal conditions: Task-Relevant AM, Task-Irrelevant AM, and Pure Tagging (Figure 2a; see Supplementary Table S1 for all pairwise comparisons). Comprehension accuracy was higher when AM tagging was task-relevant. Specifically, participants showed significantly greater accuracy in the Task-Relevant AM condition than in both the Task-Irrelevant AM condition (t(39) = 3.15, p = 0.031, d_z_ = 0.50) and the Pure Tagging condition (t(39) = 3.14, p = 0.032, d_z_ = 0.50), with both comparisons reflecting medium effect sizes. No significant difference was observed between the Task-Irrelevant AM and Pure Tagging conditions (p > 0.05; Figure 2a). Here d_z_ reflects the standardised effect size calculated using the difference. Together, these findings indicate that behaviourally relevant amplitude modulation selectively improved speech comprehension relative to the comparison conditions.

We next examined whether response times differed across tagging conditions. Response time was defined as the latency between the presentation of the comprehension question and the participant’s response in seconds. Response times also differed significantly across conditions, as revealed by a one-way repeated-measures ANOVA (F(4,156) = 14.60, p < 0.001). Participants responded more slowly in the Task-Irrelevant AM condition than in both the Task-Relevant AM condition (t(39) = 5.52, p < 0.001, d_z_ = 0.87) and the Pure Tagging condition (t(39) = 3.58, p = 0.009, d_z_ = 0.57). This suggests that task-irrelevant visual amplitude modulation increased processing demands. These differences reflected a large effect for the comparison with Task-Relevant AM and a moderate-to-large effect for the comparison with Pure Tagging. No significant difference was observed between the Task-Relevant AM and Pure Tagging conditions after Bonferroni correction (p > 0.05) (Figure 2b).

Together, these findings show that visual amplitude modulation benefited performance only when it carried information relevant to the attended speech stream: task-relevant modulation improved comprehension, whereas task-irrelevant modulation slowed responses without improving accuracy.

### Task-Relevant Amplitude Modulation Enhances Visual Tagging and Attenuates Auditory Tagging Responses

To test whether the behavioural relevance of envelope modulation altered visual tagging responses, we compared 55 Hz coherence between the Task-Relevant and Task-Irrelevant AM conditions. Visual tagging coherence was significantly stronger when the visual flicker was modulated by the attended speech envelope, with three significant occipital clusters identified using cluster-based permutation statistics (all ps ≤ 0.05; Figure 3a). Furthermore, participants with larger condition differences in 55 Hz visual tagging coherence also showed larger differences in speech comprehension accuracy (Spearman’s ρ = 0.326, p = 0.04; Figure 3b).

**Fig. 3.**
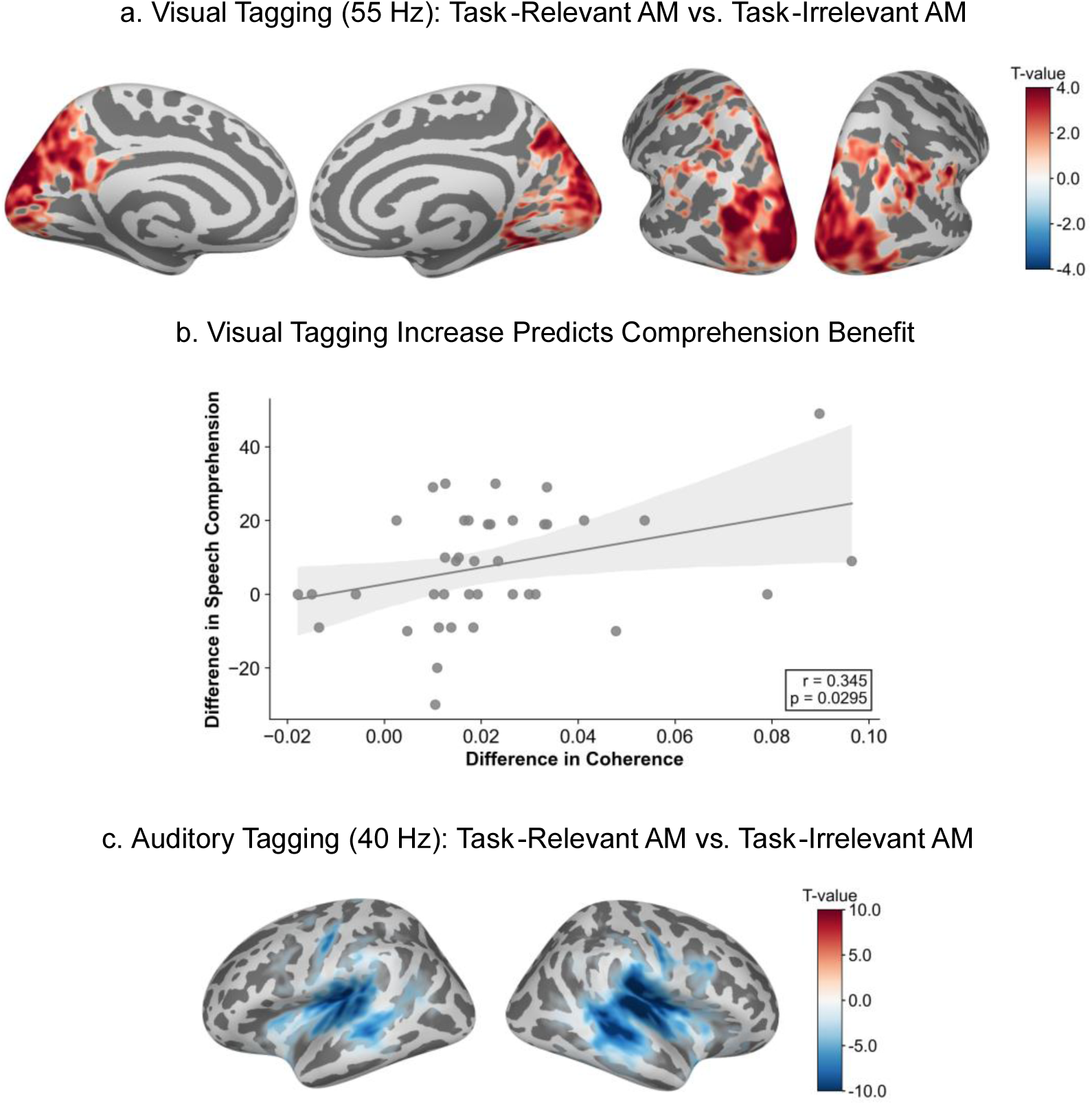
Frequency tagging responses by coherence analysis for the Task-Relevant AM vs. Task-Irrelevant AM condition. **(a)** Source-level contrast of visual tagging coherence at 55 Hz between the Task-Relevant AM and Task-Irrelevant AM conditions. Warm colours indicate significantly stronger visual tagging when the 55 Hz tagging applied to the speaker’s lip region was modulated by the attended speech envelope. **(b)** Relationship between the Task-Relevant AM vs. Task-Irrelevant AM difference in visual tagging coherence and the corresponding difference in speech comprehension accuracy. Participants showing larger increases in visual tagging coherence also exhibited greater improvements in comprehension (Spearman’s ρ = 0.326, p = 0.04). **(c)** Source-level contrast of auditory tagging coherence at 40 Hz between the Task-Relevant AM and Task-Irrelevant AM conditions. Cool colours indicate significantly stronger auditory tagging responses when the visual tagging was modulated by the task-irrelevant speech envelope. Source-level statistics were performed using a cluster-based permutation test (p < 0.05).

Auditory tagging showed the reverse pattern. Forty-hertz auditory coherence was stronger in the Task-Irrelevant AM condition than in the Task-Relevant condition across bilateral auditory and temporal regions (cluster-based permutation statistics; all ps ≤ 0.05; Figure 3c). This suggests that when visual amplitude modulation was not behaviourally relevant, participants relied more strongly on auditory speech tracking. Auditory tagging differences were not associated with differences in speech comprehension accuracy (Spearman’s ρ = 0.204, p > 0.05).

### Amplitude Modulation Alters Visual and Auditory Tagging Relative to Pure Tagging

To determine whether amplitude modulation (AM) altered sensory tagging responses, we compared the two amplitude-modulated conditions (Task-Relevant AM and Task-Irrelevant AM) with the Pure Tagging condition. Visual tagging coherence at 55 Hz was significantly lower in both AM conditions than in the Pure Tagging condition, with widespread decreases across visual regions (cluster-based permutation statistics; all ps < 0.05; Figure 4a,b; Supplementary Table S3). These findings indicate that introducing speech envelope modulation weakened steady-state visual tagging responses compared with the temporally regular flicker used in the Pure Tagging condition.

**Fig. 4.**
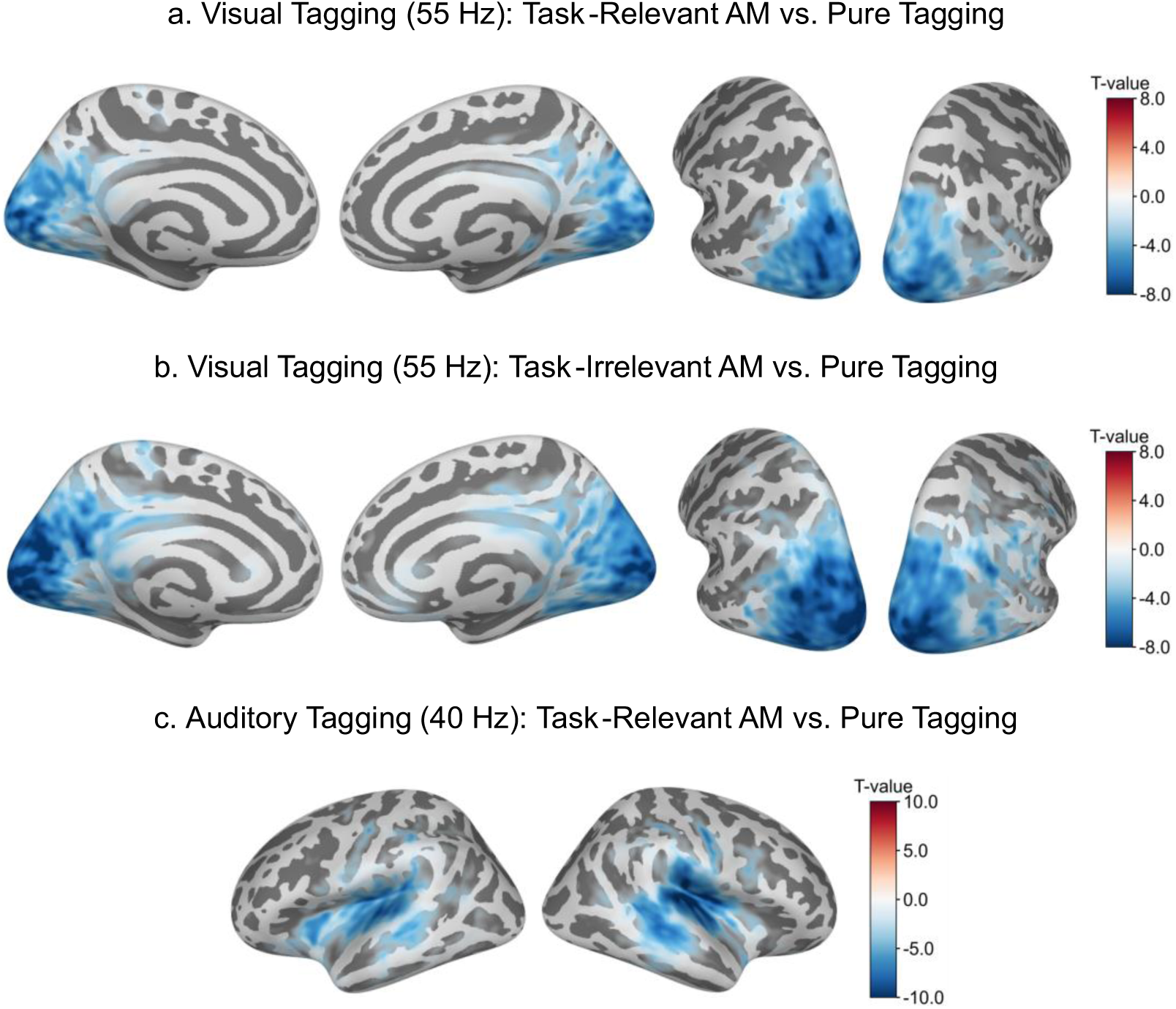
Frequency tagging responses by coherence analysis for the Task-Relevant AM vs. Pure Tagging condition. **(a)** Source-level contrast of visual tagging coherence at 55 Hz between the Task-Relevant AM and Pure Tagging conditions. Cool colours indicate significantly reduced visual tagging coherence in the Task-Relevant AM condition. **(b)** Source-level contrast of visual tagging coherence at 55 Hz between the Task-Irrelevant AM and Pure Tagging conditions. Likewise, it reveals widespread reductions in visual tagging coherence relative to Pure Tagging. **(c)** Source-level contrast of auditory tagging coherence at 40 Hz between the Task-Relevant AM and Pure Tagging conditions. Cool colours indicate significantly reduced auditory tagging coherence in the Task-Relevant AM condition. No significant auditory differences were observed between the Task-Irrelevant AM and Pure Tagging conditions. Source-level statistics were performed using a cluster-based permutation test (p < 0.05).

Amplitude modulation of visual tagging frequency had a more selective effect on auditory tagging. Relative to Pure Tagging, the Task-Relevant AM condition showed significantly reduced 40 Hz auditory coherence with a large cluster spanning auditory regions (cluster-based permutation statistics; all ps < 0.05; Figure 4c; Supplementary Table S3). No significant auditory-coherence difference was observed between the Task-Irrelevant AM and Pure Tagging conditions. This suggests that auditory tagging was reduced specifically when visual amplitude modulation carried behaviourally relevant speech envelope information, rather than in the absence of speech envelope modulation.

Together, these findings reveal two complementary effects of amplitude modulation. First, relative to Pure Tagging, speech envelope modulation generally reduced 55 Hz visual tagging coherence, likely reflecting the disruption of steady-state visual entrainment by slower luminance fluctuations. Second, within the amplitude-modulated conditions, task relevance shifted the balance between visual and auditory tracking: behaviourally relevant visual modulation enhanced visual tagging while attenuating auditory tagging. This opposing pattern suggests that audiovisual speech processing was reweighted according to task relevance, with behaviourally relevant visual speech envelope information enhancing visual speech tracking while reducing auditory tagging.

### Task-Relevant Visual Modulation Enhances Audiovisual Intermodulation Linked to Speech Comprehension

To assess nonlinear audiovisual interactions, we examined coherence at the 15 Hz intermodulation frequency, defined as the difference between the visual (55 Hz) and auditory (40 Hz) tagging frequencies. A surrogate 15 Hz intermodulation reference signal was generated by fitting sinusoids to the recorded photodiode and auditory signals around 55 Hz and 40 Hz (± 3 Hz), respectively, multiplying the fitted signals, and filtering the resulting signal to isolate the 15 Hz difference-frequency component. Coherence between neural activity and this surrogate reference signal was then compared between conditions. A cluster-based permutation comparison between Task-Relevant AM and Task-Irrelevant AM conditions revealed two significant clusters in the left motor cortex and right frontal cortex (both ps < 0.05; Figure 5a; Supplementary Table S3). Intermodulation coherence at 15 Hz was stronger in the Task-Relevant AM condition, suggesting that task-relevant visual speech envelope modulation enhanced nonlinear audiovisual interaction.

**Fig. 5.**
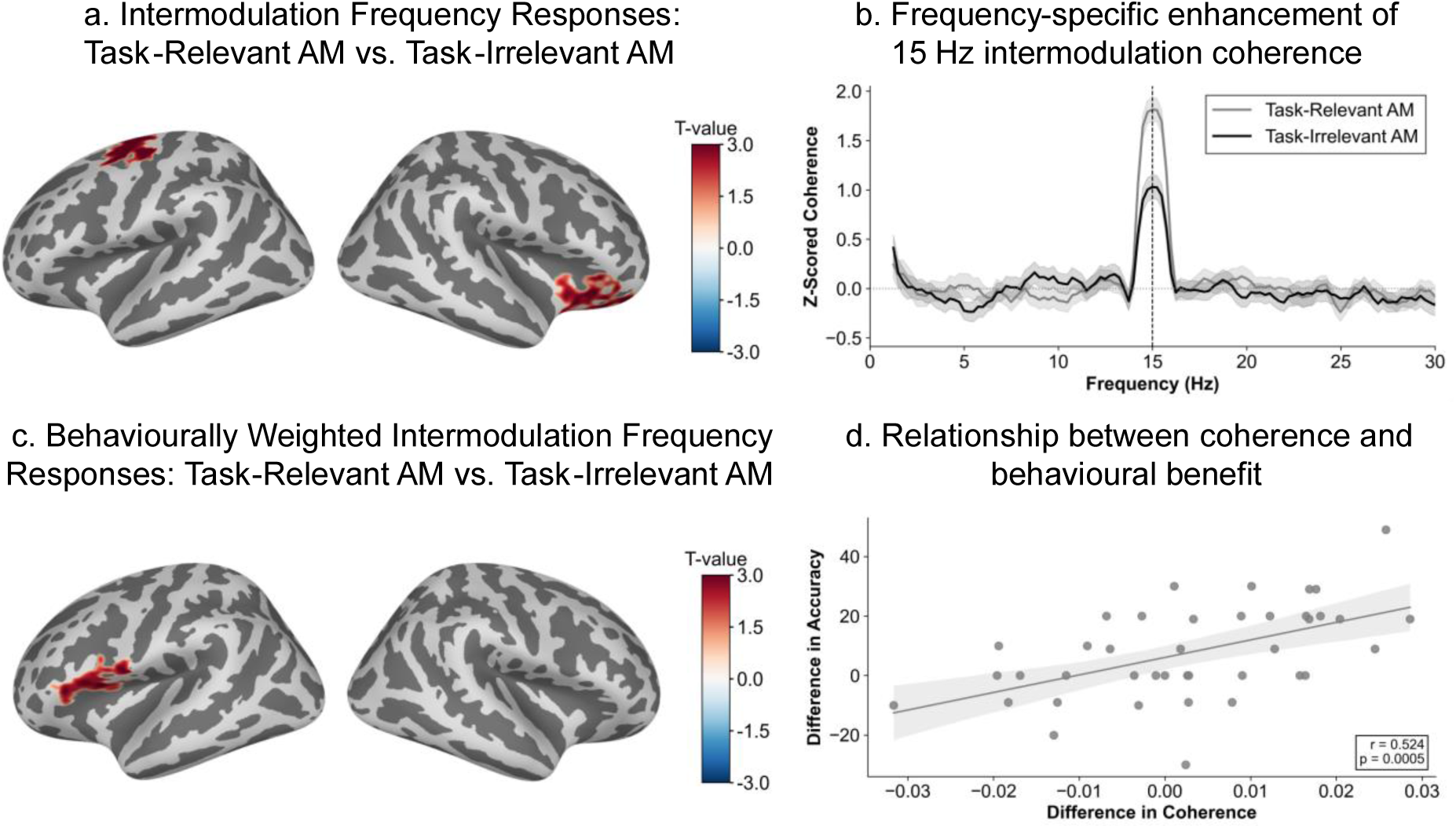
Intermodulation frequency responses by coherence analysis. **(a)** Source-level contrast of 15 Hz intermodulation coherence between the Task-Relevant AM and Task-Irrelevant AM conditions, revealing significant clusters in the left motor and right frontal regions. **(b)** Broadband coherence spectra extracted from the significant 15 Hz source-space clusters are shown as subject-wise z-scored coherence spectra and averaged across participants. Coherence was greater in the Task-Relevant AM condition than in the Task-Irrelevant AM condition at the intermodulation frequency at 15 Hz. Peaks are shown after Gaussian smoothing for visualisation. **(c)** Behaviourally weighted intermodulation analysis revealed a significant left frontal cluster in which Task-Relevant vs. Task-Irrelevant AM differences in 15 Hz intermodulation coherence covaried with corresponding differences in speech comprehension accuracy across participants. **(d)** Brain–behaviour relationship within the left inferior frontal cluster identified in (c). Mean Task-Relevant AM vs. Task-Irrelevant AM differences in 15 Hz intermodulation coherence, averaged across all vertices in the cluster, are plotted against the corresponding differences in speech comprehension accuracy. Participants with larger increases in intermodulation coherence also showed larger improvements in comprehension accuracy (Spearman’s ρ = 0.524, p < 0.001). Because the cluster was defined using the same behavioural contrast, this scatterplot is presented only to illustrate the direction of the association and does not provide an independent statistical test. Source-level statistics were performed using a cluster-based permutation test (p < 0.05).

To determine whether the condition difference was specific to the 15 Hz intermodulation frequency, rather than reflecting a broadband increase in coherence, we extracted broadband coherence spectra from the significant source-space clusters (Figure 5b). Both conditions exhibited a clear coherence peak centred on 15 Hz, corresponding to the intermodulation frequency generated by the interaction between the visual and auditory tagging. Importantly, coherence at this frequency was greater in the Task-Relevant AM condition than in the Task-Irrelevant AM condition, consistent with the source-space statistical results.

In a subsequent analysis, we tested whether condition differences in 15 Hz intermodulation coherence were related to individual differences in speech comprehension improvement. Participant-level coherence differences between the Task-Relevant AM and Task-Irrelevant AM conditions were weighted by the corresponding differences in speech comprehension accuracy before the statistical testing. This behaviourally weighted analysis identified a significant cluster in the left inferior frontal cortex (cluster-based permutation statistics; p < 0.05; Figure 5c; Supplementary Table S3), indicating that stronger task-relevance-related intermodulation responses in this region were associated with greater improvements in speech comprehension.

To illustrate this brain-behaviour relationship underlying this effect, mean coherence differences were extracted from the significant left inferior frontal cluster and correlated with speech comprehension difference scores. The results show that participants with larger increases in 15 Hz intermodulation coherence showed greater improvements in speech comprehension (Spearman’s ρ = 0.524, p < 0.001; Figure 5d). This pattern further suggests that nonlinear audiovisual interaction in the left inferior frontal cortex is linked to individual gains in speech comprehension.

## Discussion

Here we demonstrate that non-invasive visual stimulation can improve speech comprehension by embedding the attended speech envelope into naturalistic visual input. Using amplitude-modulated (AM) frequency tagging, we causally manipulated the visual speech by embedding the speech envelope of either the attended or unattended auditory stream into the speaker’s mouth region. This allowed us to test whether task-relevant visual envelope information improves speech comprehension while simultaneously measuring visual tagging, auditory tagging, and their 15 Hz intermodulation response. Behaviourally, speech comprehension improved only when the visual modulation followed the attended speech stream. At the neural level, task-relevant visual modulation enhanced visual tagging, reduced auditory tagging relative to task-irrelevant modulation, and increased 15 Hz intermodulation responses between the visual and auditory tags. Furthermore, left inferior frontal intermodulation responses covaried with individual gains in speech comprehension. Together, these findings suggest that auditory-envelope modulation of visual speech improves comprehension not simply by increasing visual salience, but by strengthening behaviourally relevant audiovisual interaction.

The behavioural results show that the benefit of visual modulation depended on its relevance to the attended speech stream. Speech comprehension accuracy was higher in the Task-Relevant AM condition than in both the Task-Irrelevant AM and Pure Tagging conditions. This distinction is important because both amplitude-modulated conditions introduced visible luminance fluctuations around the mouth, but only the modulation carrying the attended envelope improved comprehension. The effect therefore cannot be explained solely by the presence of visual flicker or by increased visual salience. Instead, the visual signal had to contain temporal information that was relevant to the speech stream participants were trying to understand. This finding extends previous work showing that visual speech cues improve intelligibility in challenging listening conditions^3,36–38^ and that enhancing the visibility of the mouth region can support speech perception under degraded listening conditions^39^. The present study goes beyond these demonstrations by showing that visual speech can be experimentally structured to carry the temporal dynamics of the attended auditory stream.

The visual tagging results support this interpretation. Visual coherence at 55 Hz was stronger when the visual flicker was modulated by the attended speech envelope than when it was modulated by the ignored speech envelope. Moreover, participants who showed larger task-relevance-related increases in visual tagging also showed larger improvements in comprehension. This suggests that the visual system tracked the tagged mouth region more strongly when its temporal structure was useful for the task. The finding extends previous RIFT work showing that visual tagging responses are sensitive to attentional relevance and stimulus clarity. Drijvers et al.^32^ reported stronger visual tagging responses when speech signals were clearer, and Zhigalov et al.^25^ showed that visual frequency-tagging responses scale with the allocation of visual attention. In the present study, however, the key manipulation was not simply whether the mouth region was visible or attended, but whether its temporal modulation matched the attended speech envelope. Visual tagging was enhanced when the tagged mouth region carried task-relevant speech information, and this enhancement was associated with better comprehension. Converging evidence from combined neural and behavioural measures suggests that visual information can support the prediction of upcoming speech in naturalistic communicative settings^40^.

Auditory tagging showed a complementary pattern. Forty-hertz auditory coherence was reduced in the Task-Relevant AM condition relative to both the Task-Irrelevant AM and Pure Tagging conditions. This reduction should not be interpreted as a loss of auditory speech processing. Rather, it suggests that auditory tracking was altered when the visual signal carried information about the attended speech stream. This interpretation is consistent with evidence that attention and task demands shape multisensory integration^41,42^, and that visual speech can modulate auditory cortical processing^43,44^. Related work has further shown that neural tracking of continuous speech is systematically influenced by acoustic degradation and listening difficulty, suggesting that changes in speech tracking responses reflect adaptive processing demands under adverse listening conditions^45^. In the present study, task-relevant visual modulation enhanced visual tagging while attenuating auditory tagging, suggesting that neural tracking did not simply increase across modalities. Instead, the balance of audiovisual tracking appeared to shift toward the visual stream when it carried useful speech envelope information. This interpretation is further supported by the relationship between the brain and speech comprehension. Visual tagging differences were associated with comprehension gains, whereas auditory tagging differences were not. Thus, the behavioural improvement appears to be more closely linked to enhanced tracking of task-relevant visual speech information than to changes in auditory tagging alone. Taken together, the opposing visual and auditory tagging effects suggest a task-dependent shift in audiovisual speech processing, in which visual speech information gained functional relevance when it carried the temporal structure of the attended speech stream.

The intermodulation findings provide clear evidence that task-relevant visual modulation affected audiovisual integration, rather than only changing modality-specific tracking. The 15 Hz intermodulation response corresponds to the difference between the 55 Hz visual tag and the 40 Hz auditory tag. Because this frequency is not present as a primary tag in either modality, it provides a frequency-specific index of nonlinear interaction between the auditory and visual neuronal signals. Task-relevant modulation increased 15 Hz intermodulation coherence, with effects observed in left motor and right frontal regions, and broadband spectra confirmed that the condition difference was concentrated at the intermodulation frequency. The left motor effect is consistent with the involvement of articulatory-motor systems in audiovisual speech perception^46^, and with evidence that frontal and motor signals can modulate auditory cortical tracking during continuous speech^14^. The right frontal effect may reflect attentional control over behaviourally relevant audiovisual information, in line with accounts of right frontal involvement in attention to behaviourally relevant stimuli^47^ and the close interaction between attention and multisensory integration^42^.

A separate analysis then tested whether this intermodulation response was related to speech comprehension improvement. Previous RIFT work has shown that audiovisual integration can be indexed by intermodulation responses, including in the left inferior frontal region^32^. The present study extends that approach by using RIFT not only to measure audiovisual interaction, but also to embed speech envelope information into naturalistic visual speech. Critically, participants who showed larger increases in 15 Hz intermodulation coherence in the Task-Relevant AM relative to the Task-Irrelevant AM condition also showed larger gains in comprehension accuracy. This brain-behaviour association was localised to the left inferior frontal cortex, a region involved in connected speech processing and temporal-frontal language networks^48–50^. This suggests that nonlinear audiovisual interaction in the left inferior frontal cortex was linked to how effectively listeners used task-relevant visual temporal information to understand the attended speech.

The present findings should be interpreted in light of several limitations. First, although the high-frequency RIFT carrier was designed to be imperceptible, the speech envelope modulation produced visible luminance fluctuations around the mouth. As a result, the behavioural improvement observed may partly reflect enhanced visual saliency rather than speech-specific temporal alignment driven by the RIFT per se. In addition, the current study did not include a control condition in which the speech envelope was applied directly to the lip region without RIFT. Including such a condition would allow future work to determine whether the behavioural effects observed here are specific to amplitude-modulated RIFT or extend more generally to visually presented speech envelope dynamics. Second, the auditory tag was perceptible as a low-frequency buzz. Behavioural performance indicates that speech comprehension was preserved, but future work could explore less perceptible auditory tagging approaches. Finally, the present sample consisted of young adults with normal hearing. Whether the same mechanism supports comprehension in older adults or listeners with hearing difficulties remains to be tested. Despite these limitations, the present findings suggest that non-invasive sensory stimulation, when structured by task-relevant speech information, can influence speech perception under challenging listening conditions.

Beyond its theoretical implications, amplitude-modulated RIFT may have potential relevance for populations that experience difficulties understanding speech in complex listening environments. Speech-in-noise perception is known to decline with age and is strongly associated with increased listening effort and reduced sensitivity to temporal speech cues^51,52^. Older adults and listeners with hearing impairments often rely more heavily on visual speech information, suggesting that cross-modal compensatory mechanisms play a critical role when auditory processing becomes unreliable^5,53^. By delivering behaviourally relevant temporal structure via the visual modality, amplitude-modulated RIFT may provide a route for supporting speech comprehension. This distinguishes the approach from conventional hearing aids, which primarily operate through amplification and noise-reduction strategies but do not directly address multisensory integration or temporal prediction mechanisms.

Moreover, visually delivered rhythmic stimulation can operate without altering the acoustic signal itself, potentially allowing RIFT-based approaches to complement rather than replace conventional assistive technologies. Hearing aids primarily improve audibility but cannot fully compensate for neural processing limitations that affect speech perception, such as reduced temporal resolution^54^. Future studies should therefore investigate whether combining visual rhythmic stimulation with conventional amplification can improve speech comprehension in listeners who obtain limited benefit from hearing aids alone.

Furthermore, this approach may also offer advantages relative to other intervention strategies. Perceptual training and speechreading programmes can improve audiovisual speech perception but typically require prolonged learning and show substantial inter-individual variability^55^. In contrast, amplitude-modulated RIFT delivers task-relevant temporal information directly through the visual signal.

Together, these considerations suggest that visually mediated rhythmic stimulation may represent a promising avenue for supporting communication in adverse listening conditions.

## Conclusion

This study demonstrates that RIFT, a non-invasive sensory stimulation technique, can improve speech comprehension when guided by task-relevant speech envelope information. By embedding the attended speech envelope into visual speech, amplitude-modulated RIFT enhanced visual tagging, altered auditory tagging, and strengthened 15 Hz intermodulation responses between auditory and visual signals. Critically, individual gains in comprehension were associated with intermodulation responses in left inferior frontal cortex, suggesting that nonlinear audiovisual interaction was linked to the effective use of task-relevant visual information. These findings establish amplitude-modulated RIFT as a non-invasive approach for both probing and shaping audiovisual speech processing in challenging listening conditions.

## Online Methods

### Participants

A total of 47 healthy adults (23 female; age range: 19-35 years old; mean age: 23.9 ± 4.1, mean ± std) participated in this study at the Centre for Human Brain Health (CHBH), University of Birmingham. Seven participants were excluded due to excessive noise, technical faults, or poor behavioural performance. Poor behavioural performance was defined a priori as comprehension accuracy below 50%. Given the three-alternative forced-choice design (33.3% chance level), this criterion ensured that the included participants performed reliably above chance. The final dataset, therefore, comprised 40 participants (20 female; age range: 19-35 years old; mean age: 23.4 ± 2.9, mean ± std).

All participants reported no history of mental and neurological illness or neurodevelopmental disorders. No participants reported taking any antipsychotic medications. All participants gave written consent prior to completing the study and were compensated for their time. All participants were right-handed, verified by the Edinburgh Handedness Inventory^56^. All participants were native English speakers and had normal or corrected-to-normal vision and normal hearing. Normal hearing was confirmed using a pure-tone threshold test (Mimi Hearing Technologies GmbH). The study was approved by the University of Birmingham STEM ethics committee (ERN_18-0226AP28A) and was conducted in accordance with the Declaration of Helsinki.

### Frequency Tagging

The stimuli were presented using the PROPixx DLP LED projector (VPixx Technologies Inc., Canada). This projector has a refresh rate of up to 1,440 Hz. To achieve this refresh rate, each frame from the stimulus PC graphics card (120 Hz) was split into quadrants, increasing the refresh rate fourfold (480 Hz). The stimuli were presented in the centre of a grey background at a resolution of 1289 x 720 pixels. The rectangular area (120 x 90 pixels) around the speaker’s lips was tagged using 55 Hz sine waves (Figure 1). There was also a small, tagged box in the bottom right corner of the screen where the photodiode was secured to measure the luminance changes on the screen (30 x 30 pixels).

Two types of frequency tagging were included in this study - pure tagging and amplitude-modulated (AM) tagging. Pure tagging consisted of a continuous imperceptible 55 Hz sinusoidal luminance modulation of the lip region with constant amplitude. Amplitude-modulated tagging was achieved by multiplying the pure tagging signal by the auditory speech amplitude envelope (see the *Stimuli, Task and Experimental Conditions* for more details). In the Task-Relevant and Task-Irrelevant AM conditions, the amplitude modulation introduced perceptible slow luminance fluctuations. The auditory speech stream was tagged at 40 Hz by multiplying the naturalistic speech waveform by a 40 Hz sinusoidal signal. Although the resulting 40 Hz auditory tag was audible as a low-frequency buzz, behavioural performance (Figure 2) indicated that it did not interfere with speech comprehension.

### Stimuli, Task, and Experimental Conditions

The stimuli used in this study were continuous audiovisual speech videos, as used and validated by Park et al.^37^. Each video was between 7 and 9 minutes in length. The content of the videos was adapted from TED talks (www.ted.com/talks/) and was recorded by a professional male speaker. In all conditions, participants watched a video recording while simultaneously hearing two different auditory speech in each ear. Participants were instructed to which ear they should pay attention to (task-relevant speech) and which they should ignore (task-irrelevant speech) at the start of each condition. Participants attended to the same ear for all conditions. To reduce potential bias associated with attending to a particular ear, participants were unaware that the attended ear remained constant. The attended ear (left or right) was counterbalanced across participants, such that the attended ear remained the same across conditions within each participant (20 paid attention to the left- and 20 to the right-side speech). Because the attended ear was counterbalanced across participants, any potential lateralisation biases were controlled at the group level.

There were five experimental conditions; however, the analysis focused on three conditions (see Supplementary Information for the other conditions): 1) In the task-relevant amplitude modulation condition (Task-Relevant AM), the visual tagging signal at 55 Hz was modulated by the envelope of the speech that the participants were instructed to attend to. 2) In the task-irrelevant amplitude modulation condition (Task-Irrelevant AM), the visual tagging signal at 55 Hz was modulated by the envelope of the speech that the participants were instructed to ignore. 3) In the no amplitude modulation condition (Pure Tagging), the visual tagging signal at 55 Hz was not modulated by either of the speech envelopes, i.e., constant amplitude over time. (Figure 1). In all conditions, the auditory speech was frequency-tagged at 40 Hz without amplitude modulation.

### Experimental Procedure

Participants were comfortably seated in the MEG with the gantry positioned at a 60° angle within a magnetically shielded room to minimise external magnetic fields.

Stimuli were presented on a screen at a fixed viewing distance of 100 cm from the participant using the Psychophysics Toolbox^57^ in MATLAB (MathWorks, Natick, MA). Participants watched five audiovisual speech videos, and the order of the conditions was randomised across participants. Following each video, participants completed a 10-question multiple-choice comprehension questionnaire about the attended speech. Accuracy and response time were measured, and each video-questionnaire block lasted approximately 10 minutes.

### Data Acquisition

Magnetoencephalographic data were acquired using a 306-sensor Elekta Neuromag TRIUX MEG system with 102 magnetometers and 204 orthogonal planar gradiometers (MEGIN, Finland) at the Centre for Human Brain Health (CHBH), University of Birmingham. During data acquisition, the data were band-pass filtered from 0.1 to 330 Hz and sampled at 1,000 Hz. The visual tagging signal was recorded using a custom-made photodiode (Aalto Neuroimaging Centre, Finland) connected to the MEG system. A Polhemus Fastrak electromagnetic digitiser system (Polhemus Inc., USA) was used to digitise the locations of three fiducial points, the left and right preauricular points and the nasion. Four head position indicator (HPI) coils were secured and digitised, two on the forehead at least 3 cm apart and two on the left and right mastoid bone. To record the participant’s head shape, at least 300 additional head points were digitised. Vertical and horizontal electrooculography (EOG) and electrocardiography (ECG) were recorded during the experiment.

Behavioural responses were recorded using one five-button response box (NAtA Technologies, USA).

### Eye-Tracking

The participant’s eye gaze was tracked using the EyeLink 1000 Plus (SR Research, Ottawa, Canada) to ensure the participant was attending to the speaker’s lip. The eye-tracker was placed on a tray table 100 cm from the participants’ eyes. The eye-tracker recorded pupil size and horizontal and vertical position of both eyes. Eye tracking data were sampled at 1,000 Hz. At the start of each condition, a five-point calibration and validation procedure was performed. Calibration and validation were accepted only if the corresponding points were within 1 visual degree in both the horizontal and vertical directions.

### Data Preprocessing

MEG data were preprocessed and analysed using MNE Python^58^ following the standard preprocessing workflow described in the FLUX pipeline^59^. Firstly, bad sensors were identified and reconstructed. Then, Signal-Space-Separation (SSS) was applied to suppress environmental artefacts^60^. For datasets exhibiting excessive noise, Spatiotemporal SSS (tSSS) was applied^61^. Independent component analysis (ICA)^62^ was used to suppress EOG and ECG artefacts. Data were resampled at 250 Hz. Continuous data were epoched into 4-second segments across the duration of the task-relevant audio, from onset to offset.

### MEG-MRI Co-Registration and Source-level Preprocessing

T1-weighted Magnetization-Prepared Rapid Gradient-Echo (MPRAGE) structural magnetic resonance image (MRI) scans were acquired from each participant using a 3-Tesla Siemens PRISMA scanner (TR = 2,000 ms, TE = 2.03 ms, TI = 880 ms, flip angle = 8°, FOV = 256 × 256 × 208 mm, 1 mm isotropic voxel). FreeSurfer recon-all was used to reconstruct surfaces. A Boundary Element Method (BEM) watershed was used to create the scalp, inner and outer skull^58^. The source space was defined based on the brain surface, then the forward solution was calculated. For each participant, the structural MRI scans were aligned with the MEG data by spatially co-registering the three fiducial points from the digitisation at the MEG session. Then the Iterative Closest Point (ICP) algorithm^63^ was used to align the digitised head shape with the MRI head surface.

### Behavioural Data Analysis

Behavioural accuracy was analysed to assess speech comprehension. Using Python, a one-way repeated-measures analysis of variance (ANOVA) was used to evaluate the overall effect of conditions. Where appropriate, follow-up paired-samples t-tests were conducted to assess pairwise differences between conditions. The significance threshold for these post hoc comparisons was Bonferroni-corrected, with statistical significance defined as p < 0.05 after correction.

### Frequency Tagging Analysis

To characterise tagging responses at the source level, dynamic statistical parametric mapping (dSPM) source estimates were computed for each epoch, and coherence was calculated between each cortical vertex and the seed channels. The visual seed channel was a photodiode attached to the display screen (see Frequency Tagging) and the auditory seed channels were recorded using the left and right soundcard signal during the auditory speech delivery. For each 4-second epoch, coherence was estimated using a multitaper approach and averaged within ± 3 Hz around 55 Hz (visual tagging) and 40 Hz (auditory tagging). Coherence values were then averaged across epochs to obtain condition-specific source maps, which were morphed to the FreeSurfer average template (fsaverage) for group-level analysis. Cluster-based permutation tests were used to assess condition differences in tagging responses, as indexed by coherence. For each comparison, coherence values at the tagging frequency were extracted and condition differences were computed at the single-subject level. These difference values were entered into a one-sample permutation test against zero across participants. Clusters were formed using the spatial adjacency derived from the fsaverage source space, with a cluster-forming threshold applied to the t-statistics. Cluster-level statistics were computed as the sum of t-values within each cluster. Statistical significance was determined using a permutation distribution generated from 1024 random sign-flips. Cluster-level p-values were obtained by comparing observed cluster statistics to this null distribution. All tests were two-tailed, and clusters with p < 0.05 were considered significant.

### Intermodulation Signal Generation and Coherence Analysis

To characterise interactions between auditory and visual frequency-tagged signals, intermodulation reference signals were generated from the recorded photodiode and auditory channels. The recorded photodiode and audio signals were band-pass filtered around the visual (55 Hz) and auditory (40 Hz) tagging frequencies, respectively (± 3 Hz). Sinusoids were then fitted to the filtered signals, and the fitted sinusoidal signals were multiplied to derive a surrogate for the intermodulation reference signals, which were then scaled to -1 to 1 and filtered between 1 and 20 Hz to retain the 15 Hz-difference-frequency component.

Source-level coherence between the intermodulation signal and reconstructed cortical activity was computed for each participant and experimental condition. Condition-specific coherence maps were transformed to the fsaverage template and entered cluster-based permutation analyses to identify cortical regions exhibiting significant condition-related differences in intermodulation coherence. Statistical significance was assessed using cluster-based one-sample permutation testing. For each condition comparison, subject-level coherence differences were calculated at every source vertex and tested against zero using 1,024 permutations. Spatial clustering was based on the fsaverage source-space adjacency matrix. Cluster-forming thresholds were derived from the t-distribution (two-tailed, α = 0.05), and clusters with p < 0.05 were considered significant.

For significant intermodulation frequency clusters, broadband coherence spectra were extracted by averaging fsaverage-morphed coherence values across all vertices within each cluster, allowing assessment of the frequency specificity of condition-related effects shown in Figure 5b.

To identify neural responses associated with behavioural benefit, a behaviourally weighted source-space analysis was additionally performed. For each participant, coherence differences between experimental conditions were multiplied by the corresponding behavioural difference score (the difference in speech comprehension accuracy), yielding a behaviourally weighted neural estimate at each source vertex. The resulting weighted maps were entered into the same cluster-based permutation framework described above to identify cortical regions in which condition-related neural changes systematically covaried with behavioural performance across participants. As a validation analysis, participant-level coherence differences were extracted from the significant clusters identified in the behaviourally weighted analysis. For each participant, coherence differences were averaged across all vertices belonging to these clusters. These values were then correlated with differences in speech comprehension scores using Spearman’s rank correlation (Figure 5d).

## Author Contributions

HP conceived the study. HP, YP and AP developed the stimulus-presentation script. CR collected, analysed the data, and drafted the manuscript under the supervision of KS and HP. All authors contributed to the interpretation of the results, revised the manuscript, and approved the final version.

## Acknowledgements

We thank the Centre for Human Brain Health (CHBH), University of Birmingham, for infrastructure and technical support. We are grateful to Jonathan Winter for MEG technical support and assistance with data acquisition. We also thank all participants for their time and engagement.

## Code Availability

The analysis scripts will be made available through a dedicated GitHub repository upon publication.

## Data Availability

The datasets generated and analysed during the current study are available from the corresponding author upon request.

