## Supplementary Information for "Augmenting Speech Comprehension using Rapid Frequency Tagging"

### Methods

*Supplementary Experimental Conditions*

The supplementary analyses included two additional control conditions. In the Naturalistic Task-Relevant AM condition, participants viewed and listened to a single audiovisual speech stream without dichotic presentation. The same auditory speech signal was presented to both ears and was congruent with the visual speech. As in the Task-Relevant AM condition, the 55 Hz visual tag was amplitude-modulated by the speech envelope, and the auditory signal was tagged at 40 Hz. This condition was included to assess comprehension under more naturalistic audiovisual listening while preserving envelope-driven visual modulation. In the No Tagging condition, neither frequency tagging nor amplitude modulation was applied to the auditory or visual speech signals. Participants listened to two speech streams in the same dichotic-listening paradigm as in the main conditions, with the visual speech matched to the attended speech stream. This condition served as a baseline for assessing the effects of frequency tagging and amplitude-modulated visual stimulation. In both supplementary conditions, participants completed the same speech comprehension task for the attended speech stream, using the same procedure as in the main conditions.

### Results

***Behavioural Results including the Naturalistic Task-Relevant AM and No-Tagging Conditions***

To provide additional context for the effects observed in the main behavioural analyses, we examined performance in two supplementary conditions: the Naturalistic Task-Relevant AM condition and the No Tagging condition (Figure S1).

A repeated-measures ANOVA including all five experimental conditions revealed significant differences in both speech comprehension accuracy ($F(4,156) = 4.82$, $p < 0.001$; Figure S1a) and response time ($F(4,156) = 14.60$, $p < 0.001$; Figure S1b). Post hoc Bonferroni-corrected comparisons indicated that speech comprehension accuracy in the Naturalistic Task-Relevant AM condition was significantly higher than in the Pure Tagging condition ($t(39) = 3.14$, $p = 0.032$, $d_z = 0.50$), whereas no significant difference was observed between the Naturalistic Task-Relevant AM and No Tagging conditions ($p > 0.05$). In addition, accuracy in the No Tagging condition was significantly lower than in both the Task-Relevant AM condition ($t(39) = 8.28$, $p < 0.001$, $d_z = 1.31$) and the Pure Tagging condition ($t(39) = 5.95$, $p < 0.001$, $d_z = 0.94$).

These findings suggest that removing visual tagging altogether is associated with poorer speech comprehension performance, whereas naturalistic modulation provides a modest benefit relative to simple visual tagging.

For response time (Figure S1b), the No Tagging condition elicited significantly slower responses than the Task-Relevant AM condition ($t(39) = 4.75$, $p < 0.001$, $d_z = 0.75$), while no significant difference was observed between the Naturalistic Task-Relevant AM and No Tagging conditions following Bonferroni correction ($p > 0.05$). Participants also responded more slowly in the Naturalistic Task-Relevant AM condition than in the Task-Relevant AM condition, although this comparison did not survive correction for multiple comparisons ($p > 0.05$). Overall, these supplementary analyses indicate that explicit task-relevant amplitude modulation yields the most favourable combination of accuracy and response speed, whereas naturalistic modulation and the absence of tagging produce intermediate levels of performance.

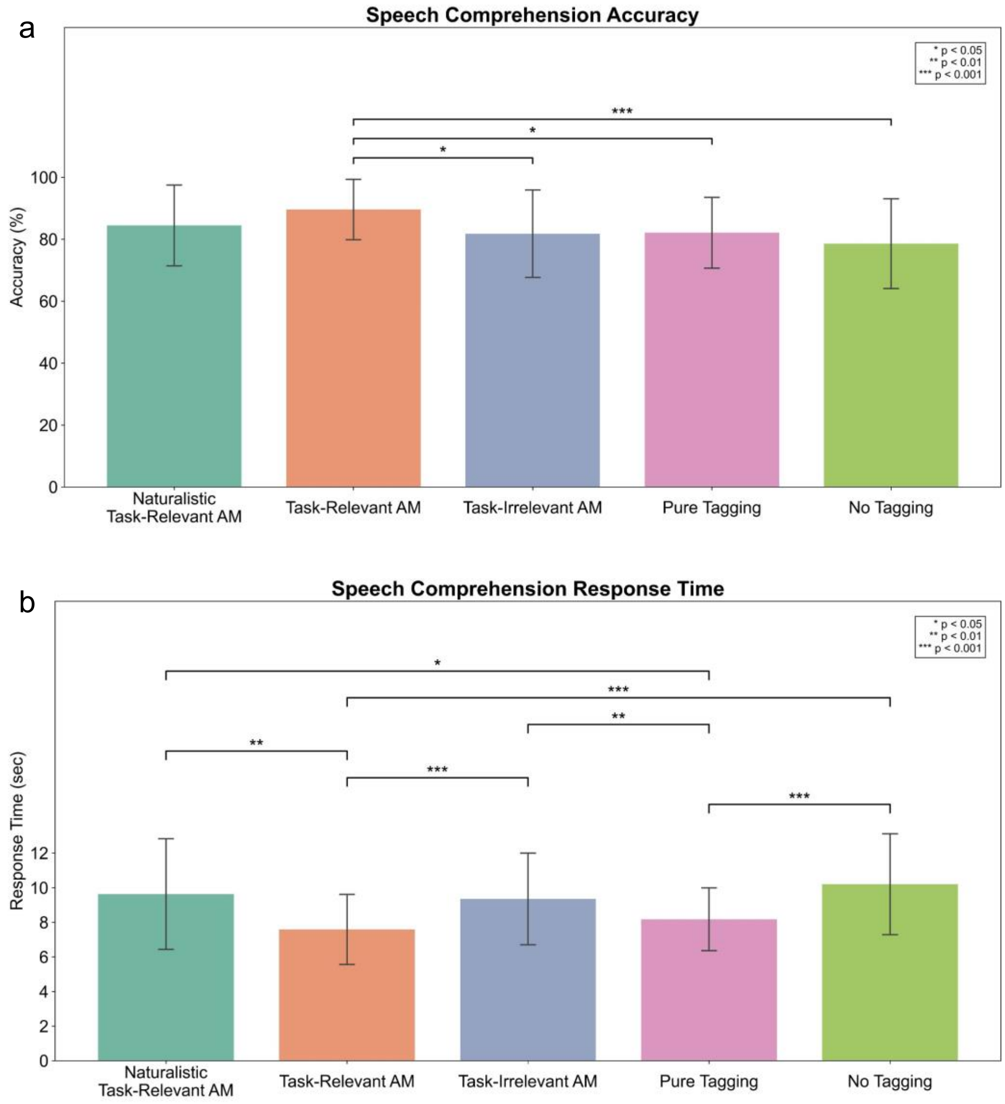


**Fig. S1 | Inclusion of Naturalistic Task-Relevant AM and No Tagging control conditions. (a)** Speech comprehension accuracy and **(b)** response time across all five experimental conditions. The supplementary Naturalistic Task-Relevant AM and No Tagging conditions provide context for the main experimental comparison between the Task-Relevant AM, Task-Irrelevant AM, and Pure Tagging conditions. Although performance in the Naturalistic Task-Relevant AM condition was intermediate between Task-Relevant AM and the control conditions,

Task-Relevant AM remained associated with the highest accuracy and fastest responses overall. Error bars represent ±1 standard deviation. Statistical significance was assessed using one-way repeated-measures ANOVAs followed by Bonferroni-corrected paired-samples t-tests.

#### *Naturalistic Task-Relevant AM Compared to Task-Relevant AM Condition*

To determine whether the neural effects observed in the main analyses were specific to selective auditory attention, we compared the Naturalistic Task-Relevant AM and Task-Relevant AM conditions. No significant differences in visual tagging coherence at 55 Hz were observed between conditions, suggesting comparable response to the envelope-modulated visual signal. In contrast, auditory tagging coherence at 40 Hz differed significantly between conditions, with clusters localised primarily to bilateral auditory regions (Figure S2). These findings suggest that auditory tagging was sensitive to whether participants listened to a single congruent audiovisual speech stream or selectively attended to one of two competing speech streams, whereas visual tagging was primarily determined by the behavioural relevance of the visual modulation.

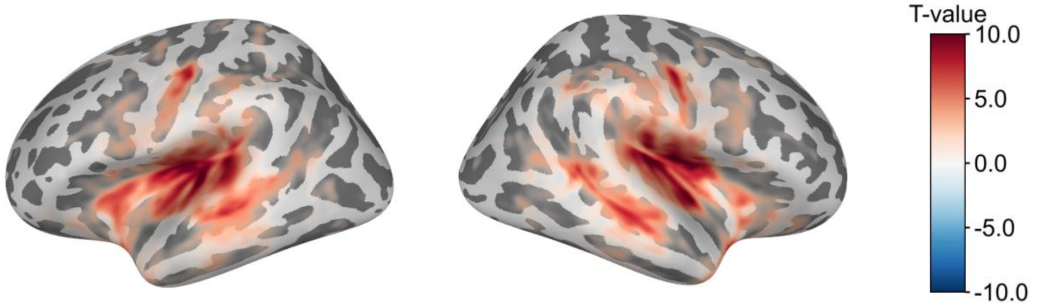


**Fig. S2 | Source-level contrast of auditory tagging coherence at 40 Hz between the Naturalistic Task-Relevant AM and Task-Relevant AM conditions.** Warm colours indicate significantly stronger auditory tagging coherence in the Naturalistic Task-Relevant AM condition relative to the Task-Relevant AM condition. No significant differences were observed for visual tagging coherence at 55 Hz. These findings suggest that auditory, but not visual, tagging is influenced by the broader listening context when visual modulation remains behaviourally relevant (cluster-based permutation statistics; $p < 0.05$).

#### *Visual Tagging Response: Tagged Conditions Compared to No Tagging Condition*

To confirm that the visual frequency tagging manipulation successfully elicited neural responses, visual tagging conditions were compared against the No Tagging condition. All tagging conditions showed significantly stronger visual tagging coherence than the No Tagging condition (all cluster-corrected ps < 0.001). Significant effects were observed bilaterally across visual cortical regions. The spatial distribution of these effects was highly consistent across conditions, confirming that both amplitude-modulated and non-modulated tagging reliably induced measurable neural tagging responses.

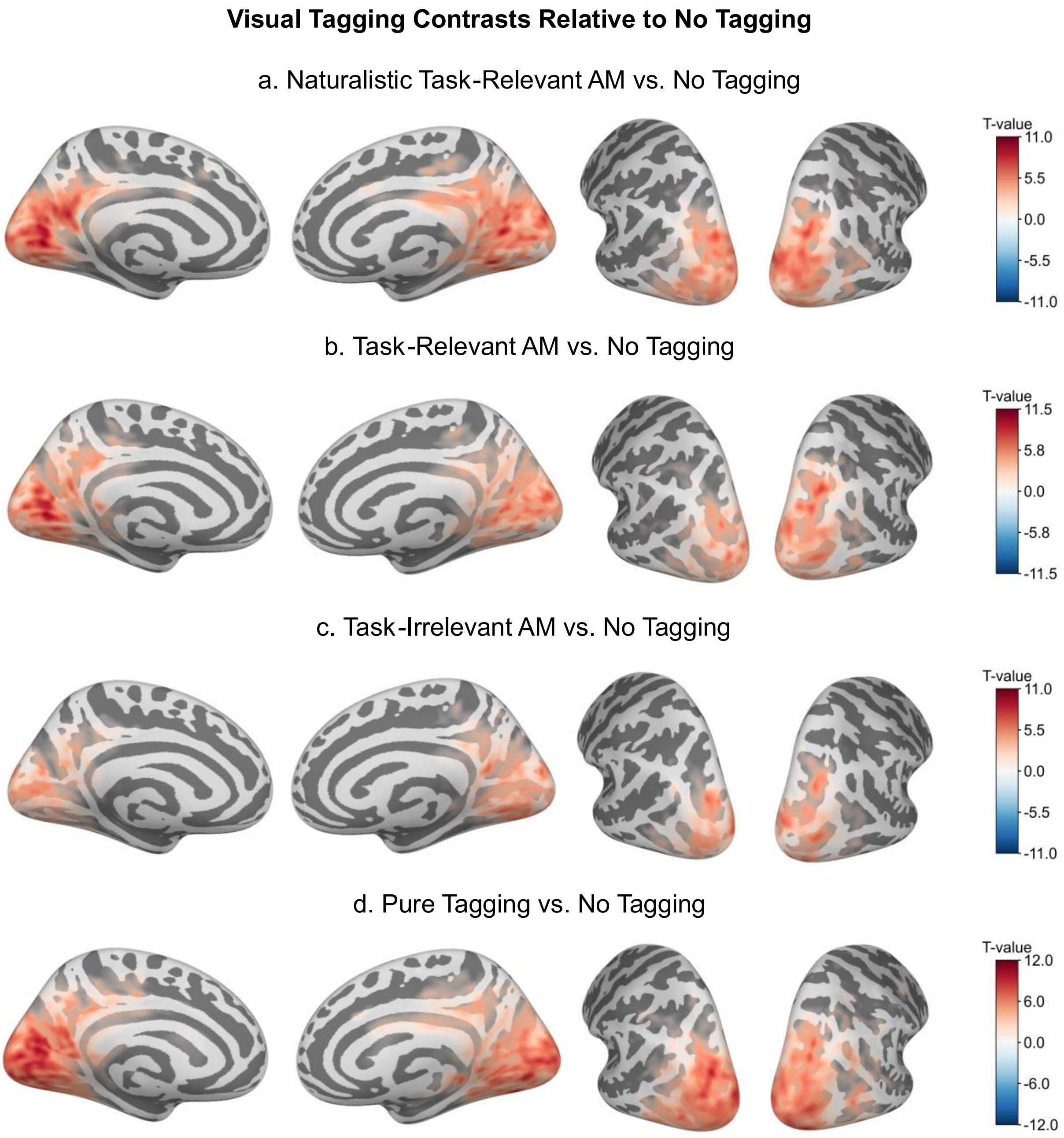


**Fig. S3 | Visual tagging responses for all tagging conditions relative to the No Tagging condition.** Source-level contrasts of visual tagging coherence at 55 Hz between each tagging condition and the No Tagging condition. **(a)** Naturalistic Task-Relevant AM vs. No Tagging, **(b)** Task-Relevant AM vs. No Tagging, **(c)** Task-Irrelevant AM vs. No Tagging, and **(d)** Pure Tagging vs. No Tagging. Warm colours indicate significantly greater visual tagging coherence in the tagging condition relative to the No Tagging condition. Colours represent cluster-corrected t-values, with only significant vertices displayed (cluster-based permutation statistics; $p < 0.05$).

### *Auditory Tagging Response: Tagged Conditions Compared to No Tagging Condition*

To confirm successful auditory frequency tagging, auditory tagging coherence at 40 Hz was compared against the No Tagging condition. All tagging conditions showed significantly stronger auditory tagging coherence than the No Tagging condition (all cluster-corrected ps < 0.001). Significant effects were observed bilaterally within auditory cortical regions, demonstrating robust entrainment of the auditory cortex by the frequency-tagged speech signal. The consistency of these effects across all tagging conditions confirms that the auditory tagging manipulation reliably elicited measurable neural responses irrespective of the accompanying visual stimulation.

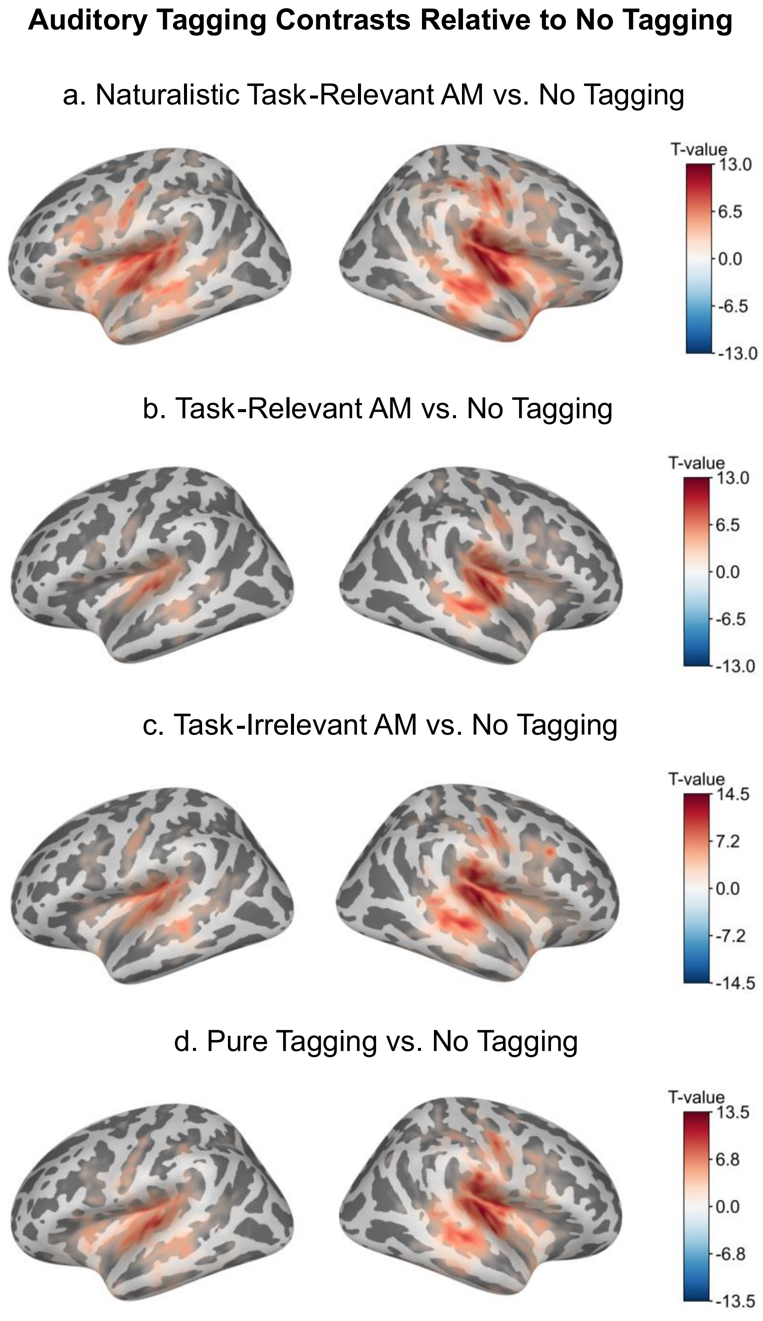


**Fig. S4 | Auditory tagging responses for all tagging conditions relative to the No Tagging condition.** Source-level contrasts of auditory tagging coherence at 40 Hz between each tagging condition and the No Tagging condition. **(a)** Naturalistic Task-Relevant AM vs. No Tagging, **(b)** Task-Relevant AM vs. No Tagging, **(c)** Task-Irrelevant AM vs. No Tagging, and **(d)** Pure Tagging vs. No Tagging. Warm colours indicate significantly greater auditory tagging coherence in the tagging condition relative to the No Tagging condition. Colours represent cluster-corrected t-values, with only significant vertices displayed (cluster-based permutation statistics; $p < 0.05$).

### *Complete Behavioural Statistics*

#### Table S1. Accuracy: Significant Bonferroni-Corrected Pairwise Comparisons

Repeated measures ANOVA: $F(4,156) = 4.82$, $p < 0.001$

| Comparison | t(39) | pBonf | Cohen's $d_z$ |
|---|---|---|---|
| No Tagging vs. Task-Relevant AM | 4.75 | < 0.001 | 0.75 |
| Pure Tagging vs. Task-Relevant AM | 3.14 | 0.032 | 0.50 |
| Task-Irrelevant AM vs. Task-Relevant AM | 3.15 | 0.031 | 0.50 |

Note. Only comparisons surviving Bonferroni correction are shown. Cohen's $d_z$ was calculated for paired-samples comparisons as $d_z = t/\sqrt{n}$, where n = 40.

#### Table S2. Response Time: Significant Bonferroni-Corrected Pairwise Comparisons

Repeated measures ANOVA: $F(4,156) = 14.60$, $p < 0.001$

| Comparison | t(39) | pBonf | Cohen's $d_z$ |
|---|---|---|---|
| Naturalistic Task-Relevant AM vs. Pure Tagging | 3.14 | 0.032 | 0.50 |
| Naturalistic Task-Relevant AM vs. Task-Relevant AM | 4.22 | 0.001 | 0.67 |
| No Tagging vs. Pure Tagging | 5.95 | < 0.001 | 0.94 |
| No Tagging vs. Task-Relevant AM | 8.28 | < 0.001 | 1.31 |
| Pure Tagging vs. Task-Irrelevant AM | 3.58 | 0.009 | 0.57 |
| Task-Irrelevant AM vs. Task-Relevant AM | 5.52 | < 0.001 | 0.87 |

Note. Only comparisons surviving Bonferroni correction are shown. Cohen's $d_z$ was calculated for paired-samples comparisons as $d_z = t/\sqrt{n}$, where n = 40.

### *Complete MEG Cluster Statistics*

#### Table S3. Summary of Significant Source-level MEG Cluster Statistics

| Comparison | Measure | Tagging Frequency | Significant Clusters (n) | Cluster p-value(s) |
|---|---|---|---|---|
| Naturalistic Task-Relevant AM vs. Task-Relevant AM | Auditory tagging coherence | 40 Hz | 2 | 0.001, 0.001 |
| Task-Relevant AM vs. Task-Irrelevant AM | Visual tagging coherence | 55 Hz | 3 | 0.010, 0.001, 0.001 |
| Task-Relevant AM vs. Task-Irrelevant AM | Auditory tagging coherence | 40 Hz | 2 | 0.001, 0.001 |
| Task-Relevant AM vs. Pure Tagging | Visual tagging coherence | 55 Hz | 2 | 0.001, 0.001 |
| Task-Relevant AM vs. Pure Tagging | Auditory tagging coherence | 40 Hz | 2 | 0.001, 0.001 |

| | | | | |
|---|---|---|---|---|
| Task-Irrelevant AM vs. Pure Tagging | Visual tagging coherence | 55 Hz | 2 | 0.001, 0.001 |
| Task-Relevant AM vs. Task-Irrelevant AM | Audiovisual intermodulation coherence | 15 Hz | 2 | 0.002, 0.012 |
| Task-Relevant AM vs. Task-Irrelevant AM | Behaviour-weighted audiovisual intermodulation coherence | 15 Hz | 1 | 0.041 |
| Naturalistic Task-Relevant AM vs. No Tagging | Visual tagging coherence | 55 Hz | 2 | 0.001, 0.001 |
| Task-Relevant AM vs. No Tagging | Visual tagging coherence | 55 Hz | 2 | 0.001, 0.001 |
| Task-Irrelevant AM vs. No Tagging | Visual tagging coherence | 55 Hz | 2 | 0.001, 0.001 |
| Pure Tagging vs. No Tagging | Visual tagging coherence | 55 Hz | 2 | 0.001, 0.001 |
| Naturalistic Task-Relevant AM vs. No Tagging | Auditory tagging coherence | 40 Hz | 2 | 0.001, 0.001 |
| Task-Relevant AM vs. No Tagging | Auditory tagging coherence | 40 Hz | 2 | 0.001, 0.001 |
| Task-Irrelevant AM vs. No Tagging | Auditory tagging coherence | 40 Hz | 2 | 0.001, 0.001 |
| Pure Tagging vs. No Tagging | Auditory tagging coherence | 40 Hz | 2 | 0.001, 0.001 |

Statistical significance was assessed using cluster-based permutation testing. Only analyses yielding significant clusters are reported. Multiple significant clusters within a comparison are listed by their cluster-corrected p-values.
